# Deep brain stimulation device-specific artefacts in MEG recordings

**DOI:** 10.1101/2023.10.25.563956

**Authors:** Bahne H. Bahners, Roxanne Lofredi, Tilmann Sander, Alfons Schnitzler, Andrea A. Kühn, Esther Florin

## Abstract

**Background:** Deep brain stimulation (DBS) has strong beneficial effects for treating movement disorders. The related cortical mechanisms can be studied with magnetoencephalography (MEG) during active DBS. However, MEG is prone to artefacts induced by the electrical stimulation and the movement of ferromagnetic DBS components. Although artefacts might vary between DBS devices from different manufacturers, no such comparison has been performed. To date, no combined MEG-DBS studies have been conducted within Yokogawa MEG systems.

**Objective:** The aim of the present study was to compare DBS artefacts in MEG phantom recordings acquired with two MEG systems (Neuromag, Yokogawa) using DBS devices from three different manufacturers (Abbott, Boston Scientifc, Medtronic) and to test whether established cleaning methods can sufficiently reduce artefacts.

**Methods:** DBS devices, electrodes, and extension cables were attached to a gelatine MEG phantom, and data was acquired in a Neuromag and a Yokogawa MEG system.

**Results:** There are device-specific differences in movement-related artefacts with weaker artefacts for Boston Scientific (BSC) devices. Stimulation-, movement- and IPG-related artefacts are best cleaned when combining the ICA-MI and Hampel filter across recordings and DBS devices. However, the cleaning of movement-related artefacts can result in artefactual spectral peaks in physiologically relevant frequencies below 20 Hz.

**Conclusions:** Device-specific IPG-related artefacts have to be considered for MEG-DBS studies and can be cleaned with combinations of published cleaning methods for MEG and EEG data. Critically, cleaning movement-related artefacts can potentially result in spectral peaks, which resemble physiological activity. Finally, combined MEG-DBS recordings are feasible in Yokogawa-MEG systems.

## Introduction

Deep brain stimulation (DBS) is a highly effective treatment in therapy refractory movement disorders (Deuschl et al., 2006; Krack et al., 2019). However, its mechanisms of action are still elusive, and the proposed concepts are under debate (Chiken and Nambu, 2016; Lozano et al., 2019). The electrophysiological investigation of DBS mechanisms started as early as the first clinical DBS trials and is an ongoing field of research (Benabid et al., 1994; Harmsen et al., 2018; Limousin et al., 1998b, 1998a; Litvak et al., 2021). Using magnetoencephalography (MEG), noninvasive recordings of cortical neural activity can be performed during active DBS. However, MEG is prone to artefacts induced by electrical stimulation and movement of ferromagnetic DBS hardware components (Bahners et al., 2020; Kandemir et al., 2020; Litvak et al., 2010; Oswal et al., 2016b).

The electrical DBS artefact results from the pulse-shaped stimulation consisting of narrow square waves (60 to 90μs), which have infinite odd harmonics of their fundamental frequency (Oppenheim et al., 1997). In theory, digitizing the electrical DBS artefact would require infinite sampling rates (Kandemir et al., 2020). Under-sampled or missed pulses result in peaks across the frequency spectrum (Jech et al., 2006; Sun et al., 2014). Artefacts induced by the movement of ferromagnetic DBS hardware, i.e. the impulse generator (IPG), extension cables, and connectors contaminate the recording at lower frequencies (Bahners et al., 2020; Kandemir et al., 2020). These components move due to the arterial pulsation and breathing and create artefacts on channels on the side of IPG implantation (Bahners et al., 2020; Litvak et al., 2010).

To reduce DBS artefacts in MEG recordings several methods have been proposed and tested systematically in phantom recordings (Kandemir et al., 2020). Especially, the independent component analysis-mutual information (ICA-MI) approach and temporal signal space separation (tSSS) are promising at the sensor and source-level (Abbasi et al., 2016; Bahners et al., 2023; Kandemir et al., 2020; Spooner et al., 2023; Taulu and Hari, 2009).

The standard protocol for DBS studies is to compare DBS on and off. Therefore, phantom recordings with different IPGs turned on and off are needed to identify potential differences of DBS artefacts. Additionally, no systematic comparison between artefacts across DBS systems from different manufacturers exists, even though hardware and software characteristics may differ between devices. Furthermore, no DBS-MEG study has been performed in a Yokogawa (YKG) MEG system. The aim of the present study was thus to characterize DBS artefacts based on phantom recordings with three different DBS devices in two MEG systems (NMG and YKG) and to test whether established artefact reduction methods can sufficiently reduce artefacts to study oscillatory activity during active DBS.

## Material and methods

### Phantom

To cast a gelatine phantom for MEG recordings, we built a mold from a commercially-available mannequin head. The gelatine powder (GoGun Ballistic 3 Gelatine, GoGun GmbH, Essen, Germany) was solved in saline solution, glycerin and isopropanol (to ensure durability) and heated to approximately 40°C. Then the gelatine was poured into the mold to allow it to cool and solidify. Next, two trajectory holes were pierced into the gelatine to later place the different DBS electrodes (Fig. 1 A). We used DBS impulse generators (IPG), extension cables and electrodes provided by three different manufacturers: Abbott (ABT, Abbott Laboratories, Abbott Park, IL, USA): Infinity 7 (model 6662), DBS electrodes (model 6170, 0.5 mm spacing) and extension cables (model 6372 and 6373); Boston Scientific (BSC, Boston Scientific Corporation, Marlborough, MA, USA): Vercise PC (model DB-1140), DBS electrodes (Vercise Cartesia, model DB-2202-30, 0.5 mm spacing) and extension cables (model NM-3138-55); Medtronic (MDT, Medtronic GmbH, Meerbusch, Germany): Activa PC (model 37601), DBS electrodes (model 3387, 1.5 mm spacing) and extension cables (model 37086). Each of the electrode pairs was inserted into the phantom head and connected to the respective IPG via extension cables. The cables were placed in one loop on the right side on top of the phantom head. From there the extensions went along the phantom’s neck to the IPG. The IPG was placed on the phantom’s right shoulder region (Fig. 1 B). All of the hardware was attached to the phantom using medical tape (Fixomull, BSN medical, Hamburg, Germany). To simulate brain signal a signal generator (PicoScope 2204A, PicoTechnology, St Neots, UK) produced a 13 Hz sinusoid within two electrode contacts of an additional discarded Medtronic DBS electrode (model 3387). The ground wire of the signal generator was connected to the ground of the magnetically-shielded room (Fig. 1 A). We inserted the electrode into a hole of 1.5 cm depth, that was located approximately 2.5 cm left from the sagittal midline and 2.0 cm behind a frontal line, matching the 10-20 EEG system frontal electrodes.

**Figure 1.**
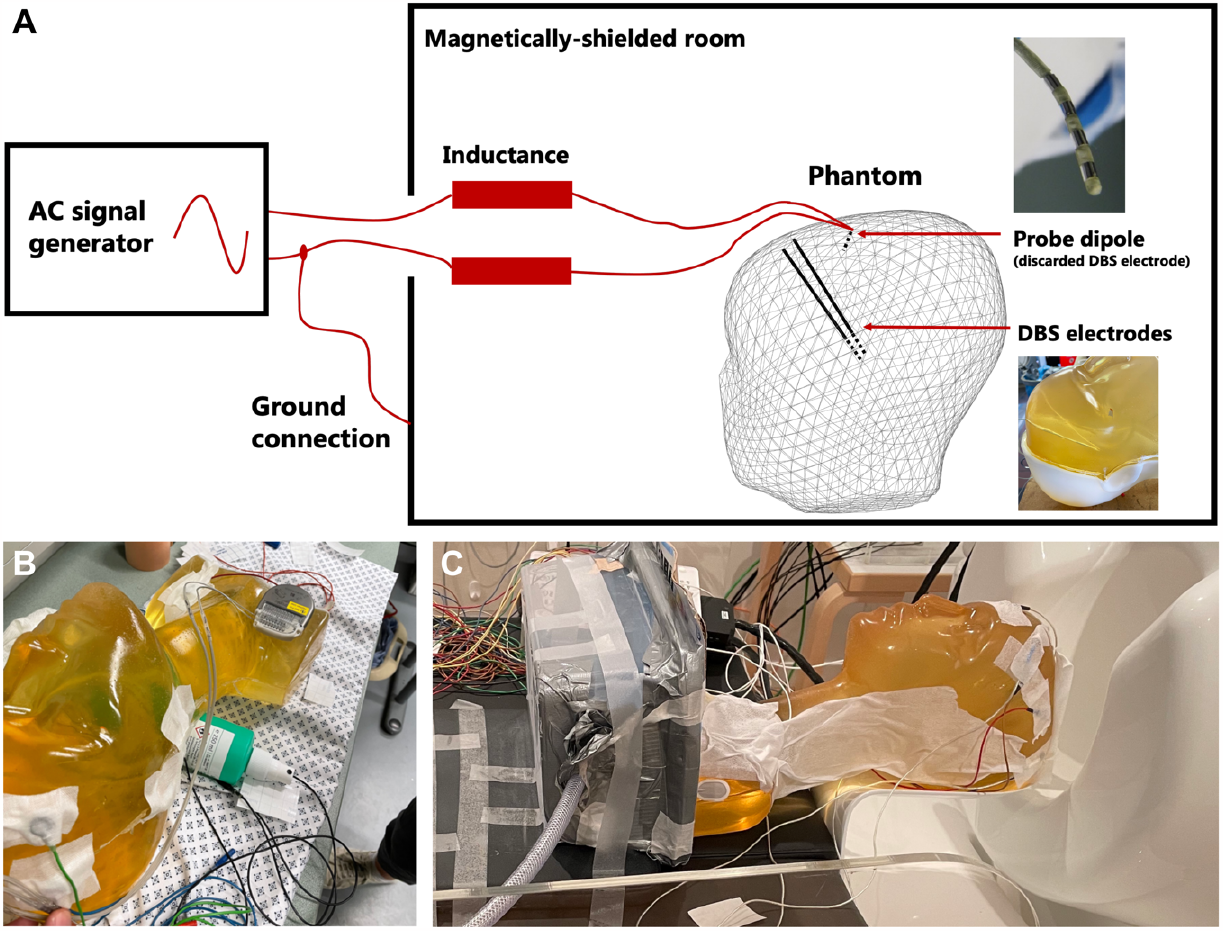
Phantom setup and experimental apparatus. **A**. The AC signal generator produced a dipole signal and was attached to a discarded DBS electrode. The respective cables were grounded with the magnetically-shielded room (MSR). The Phantom head contained two DBS electrodes that were changed between the recordings. To this end the phantom was pulled out of the MEG helmet with the MEG bed, but was not moved in relation to the inflatable box (see **C**). **B**. The IPG was placed on the right chest region of the phantom and the cables were led along the neck and to the right side of the head.

### Recordings

Phantom recordings were performed in Berlin (Physikalisch-Technische Bundesanstalt, Berlin) and Düsseldorf (Institute of Clinical Neuroscience and Medical Psychology, Heinrich Heine University Düsseldorf). In Berlin, data was acquired using a 125-channel-MEG system (Yokogawa ET 160, Tokyo, Japan). In Düsseldorf data was recorded with a 306 channel-MEG system (Neuromag, Elekta Oy, Helsinki, Finland). Data was sampled in both systems at 5,000 Hz with a high-pass of 0.1 Hz and a low-pass of 1,660 Hz.

The phantom was placed onto the respective MEG bed on top of an inflatable box, which was filled with air every 2.5 s for approximately 40 ms to simulate movement (Kandemir et al., 2020). Between recording conditions (ABT, BSC, and MDT), the phantom was not moved from its position on top of the inflatable box to keep the recording conditions constant. Instead, the phantom head was pulled out of the helmet using the respective MEG bed’s mechanism (Fig. 1 C), and only the electrodes, extension cables, and DBS devices were changed. In Düsseldorf a triaxial accelerometer (Piitulainen et al., 2013) (ADXL335 iMEMS Accelerometer, Analog Devices Inc., Norwood, MA, USA) was attached to the box to record the onset and acceleration resulting from the movement. In order to characterize the different artefacts introduced by DBS, the extension cables, as well as the movement of the phantom head inside the MEG helmet, we acquired several separate recordings of 60 s each. First, we recorded with the dipole and the movement mechanism turned on (Off+mov). This simulates a patient recording with an implanted DBS system, that is turned off. Second, we acquired a 60 s recording with the stimulation, dipole, and movement turned on (On+mov). Third, we performed a reference recording with the dipole activated and without DBS hardware (Ref). This file serves to evaluate the effectiveness of cleaning algorithms tested later on. Ideally, after cleaning the artefact, the recording should resemble the reference recording. The DBS parameters were kept constant across recordings, when switched on, with a stimulation amplitude of 4.0 mA, a pulse width of 60μs, and a stimulation frequency of 130 Hz. Due to the fact, that the size of the electrical stimulation artefact is about 25 times stronger when using monopolar compared to bipolar stimulation settings, we used a bipolar stimulation montage as typically applied in MEG-DBS studies (Bahners et al., 2020; Oswal et al., 2016b).

### Data analysis

To analyze the MEG data we used Brainstorm (Tadel et al., 2011). A notch filter was applied to reduce power line noise at 50 Hz and its harmonics up to 300 Hz in both systems. First, MEG recordings were inspected for channel jumps. Second, a power spectrum was calculated using Welch’s method with 4 s Hamming windows and 50% overlap (Welch, 1967). Third, frequency spectra were visually inspected for channels with substantially higher or lower noise levels than the average channels. For comparability reasons, we removed the same number of channels from all recordings acquired using the respective system (25 channels rejected for Neuromag recordings, 18 channels rejected for Yokogawa recordings). The accelerometer recordings from Düsseldorf were averaged across movement events for all of the three DBS devices separately to assess the comparability of recordings (Supplementary Fig. 1).

For Neuromag data, we applied the system-specific predefined signal-space projectors (SSP), designed to suppress magnetic interference from outside the magnetic-shielded room. We used three different artefact reduction approaches to clean the MEG data. First, we applied tSSS, using its MNE version as implemented in Brainstorm (Taulu and Hari, 2009). In the case of tSSS, we applied the system-specific predefined SSP of the Neuromag system after running tSSS. Second, we used the ICA-MI algorithm (Abbasi et al., 2016). Unlike in previous work, we used the channel with the largest movement artefact, identified visually from the PSD (Neuromag ABT: MEG 2423; BSC: MEG2633; MDT: MEG 1433; Yokogawa ABT: AG091, BSC: AG038, MDT: AG092) as reference channel for the mutual information calculation. Finally, we applied the Hampel filter on the data cleaned with ICA-MI and tSSS (Allen, 2009). For the sake of clarity, we do not display the results of tSSS combined with the Hampel filter in the figures.

### Artefact Comparison

To assess how effective the three different cleaning approaches are, we calculated the root mean squared error (RMSE) of the logarithmic power between the cleaned recordings and a reference recording (Kandemir et al., 2020). Namely, we compared the reference recording without an IPG inside the magnetically-shielded room and just the phantom head with the 13 Hz dipole activated to the cleaned version of the recording with stimulation and movement (On+mov). The RMSE indicates how far the cleaned sensor-level data deviate from the ground truth (reference recording) (Kandemir et al., 2020). The RMSE was computed for the average power across sensors (NMG: 179; 204 gradiometers - 25 bad channels, YKG: 104; 122 channels - 18 bad channels) in the dipole frequency range (13 Hz ± 0.5 Hz), the DBS frequency (ABT: 127 Hz, BSC and MDT: 130 Hz ± 1 Hz) and in the low-frequency range (i.e. the movement-related artefact, 1-15 Hz, excluding 13 Hz ± 0.5 Hz).

### Data availability

The raw MEG data from phantom recordings is available under the following doi: https://doi.org/10.18112/openneuro.ds004738.v1.0.0

## Results

Across devices, electrical stimulation induced artefacts at the stimulation frequency (130 Hz) or slightly below (127 Hz) in the case of ABT recordings (Fig. 2 A). Additionally, the spectrum was contaminated by further frequency peaks (e.g. ABT: 22 Hz, BSC: 22 and 25 Hz, MDT: 20 Hz) (Fig. 2 B, C). The movement of ferromagnetic hardware components resulted in low-frequency activity regardless of the device. Overall, the movement artefact was least prominent in BSC recordings (Fig. 2 B, E). The RMSE for movement artefacts is larger compared to the reference condition in the YKG system, than in the case of NMG recordings (Fig. 2, 4, 5).

**Figure 2.**
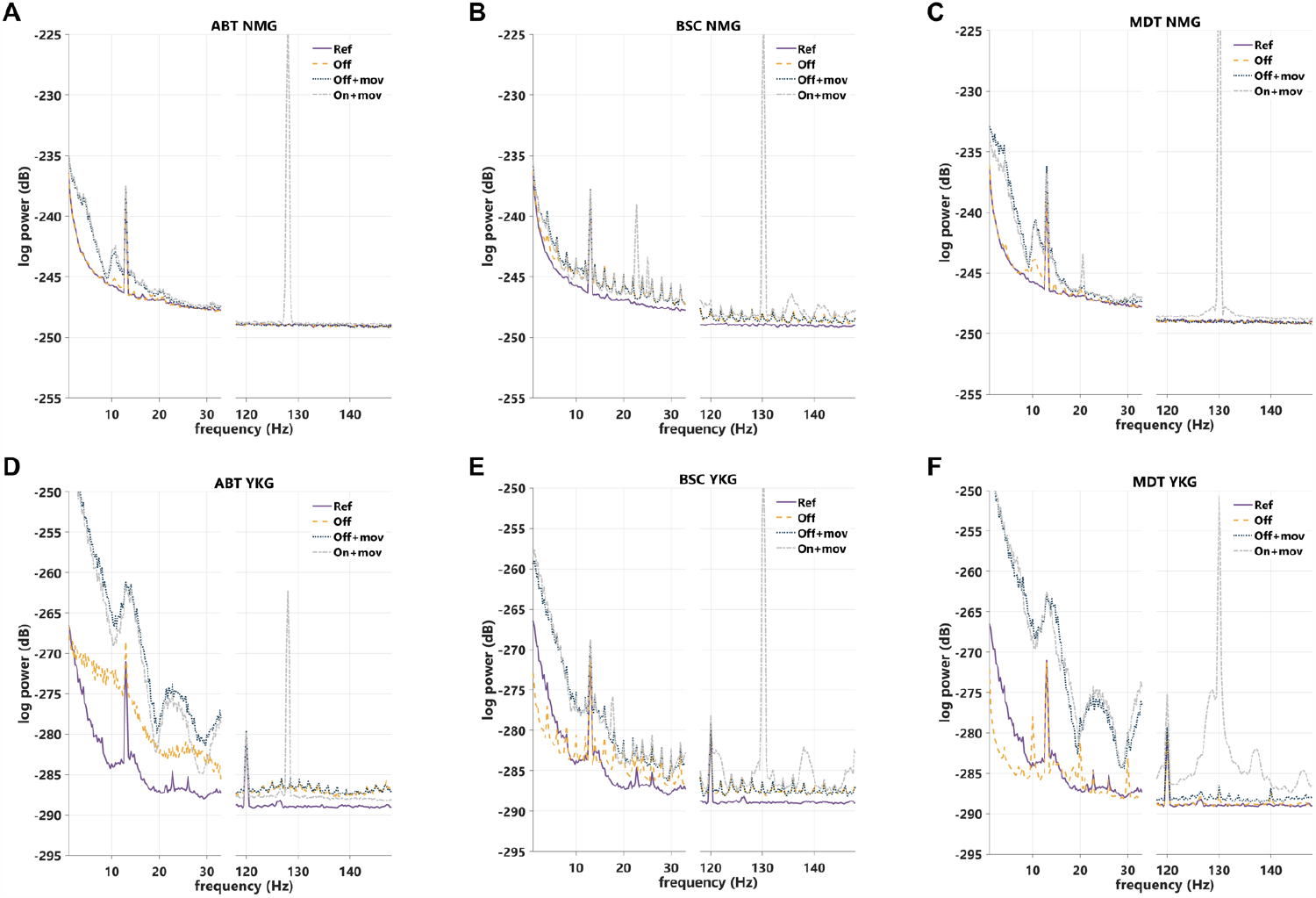
Movement-related and stimulation-induced artefacts differ across DBS devices and MEG systems. Average power spectrum density (PSD) plots across channels (gradiometers) for Neuromag (NMG) and Yokogawa (YKG) recordings (NMG: A-C, YKG: D-F). The PSD for the three DBS devices from Abbott (ABT) are shown in A and D, from Boston Scientific (BSC) in B and E and from Medtronic (MDT) in C and F. Reference recording (Ref) was performed without an implantable pulse generator (IPG) and only the dipole turned on. The Off condition was recorded with the IPG inside the magnetic-shielded room and the IPG turned Off, while only dipole activity was turned on as in the reference file. Off+mov= dipole and movement turned on and IPG turned off, On+mov= dipole, movement and IPG turned on.

Interestingly, even when switched Off, the IPGs emitted signals that led to relevant artefacts. For the BSC device, this artefact consisted of peaks with a distance of 2 Hz contaminating the spectrum in the stimulation ON and OFF condition (Fig. 2 B, E), which in the time domain appear as pulses every 100 to 500 ms (Fig. 3 E,F, Supplementary Fig. 2 E,F). In the case of the ABT device, similar artefacts with low-frequency peaks across the spectrum and an elevated low-frequency noise level occurred (Fig. 2 D, Fig. 3 A). In the time domain pulses occur about every 2 seconds and there are additional smaller pulses every 500 ms. The MDT recording showed frequency peaks with a distance of 10 Hz when the stimulation was turned off (Fig. 3 G, Supplementary Fig. 2 H). Note that IPG-related artefacts in ABT and MDT recordings were more prominent for the YKG recordings. When comparing the artefacts between DBS devices with stimulation turned on, we observe a similar overall pattern of repetitive pulses. Interestingly the pulses from the off stimulation condition disappear for the ABT and MDT device (Fig. 3 B, C, H, I), but not for the BSC system in the stimulation on condition (Fig. 3 E, F). In the case of MDT there is an additional stimulation pulse between the two 130 Hz pulses (Fig. 3 H, I). In the ABT recording, only one pulse remains in the stimulation on condition (Fig. 3 C, Supplementary Fig. 2 C), that occurs within intervals of 5 seconds.

**Figure 3.**
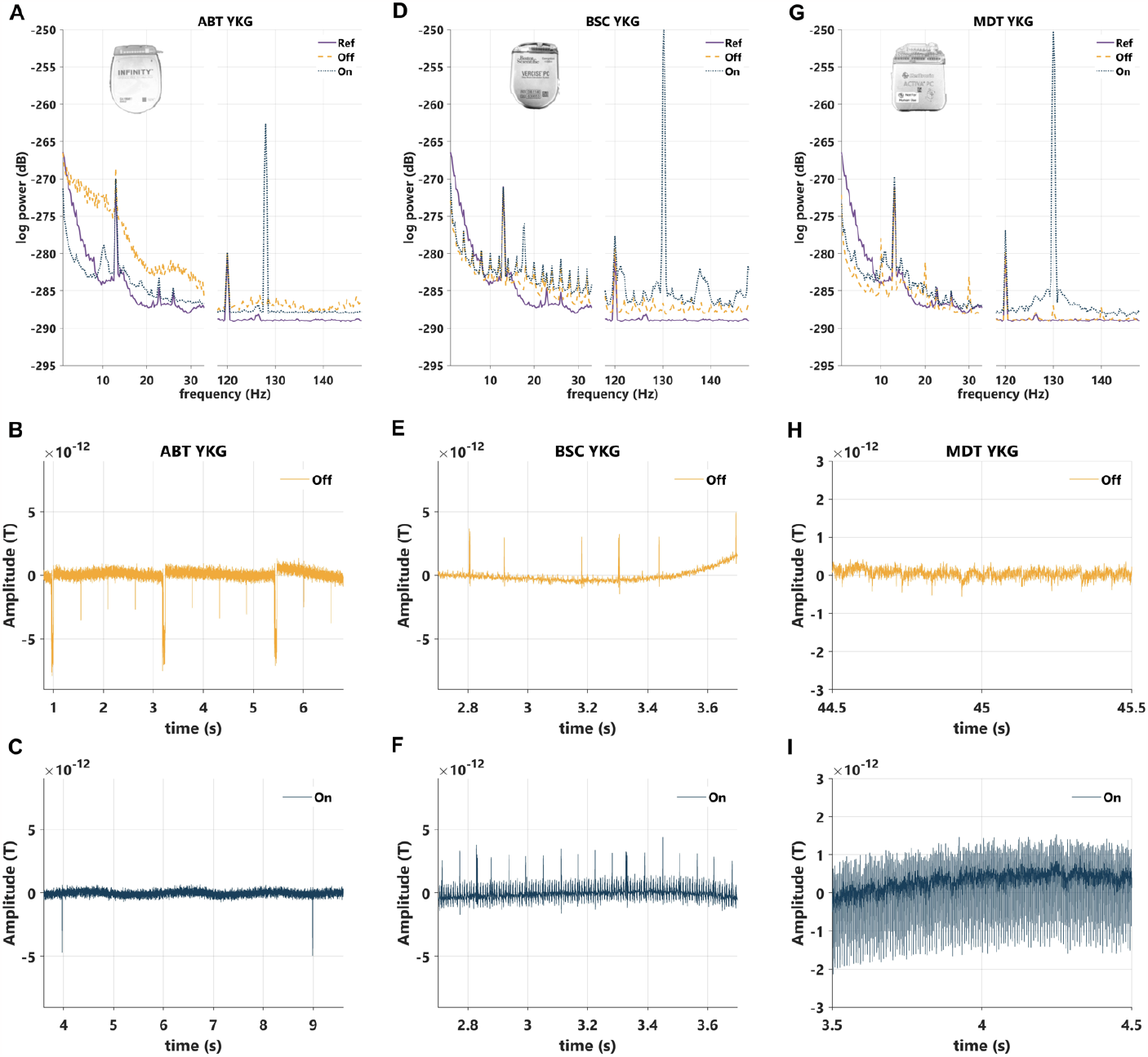
IPG-related Artefacts influence signal quality in frequency and time domain in Yokogawa recordings. **A, D, G** Average power spectrum density (PSD) plots across channels (gradiometers) for Yokogawa (YKG) recordings with DBS devices from Abbott (ABT), Boston Scientific (BSC) and Medtronic (MDT). Reference recording (Ref) was performed without an implantable pulse generator (IPG) and only the dipole turned on. The Off condition was recorded with the IPG inside the magnetic-shielded room and the IPG turned Off, while only dipole activity was turned on as in the reference file. The On condition was recorded with the IPG turned On and only the dipole activity turned on without movement. **B, C, E, F, H, I**. Exemplary time series signal showing the IPG-related artefacts in the stimulation Off (B, E, H) and On (C, F, I) conditions.

**Figure 4.**
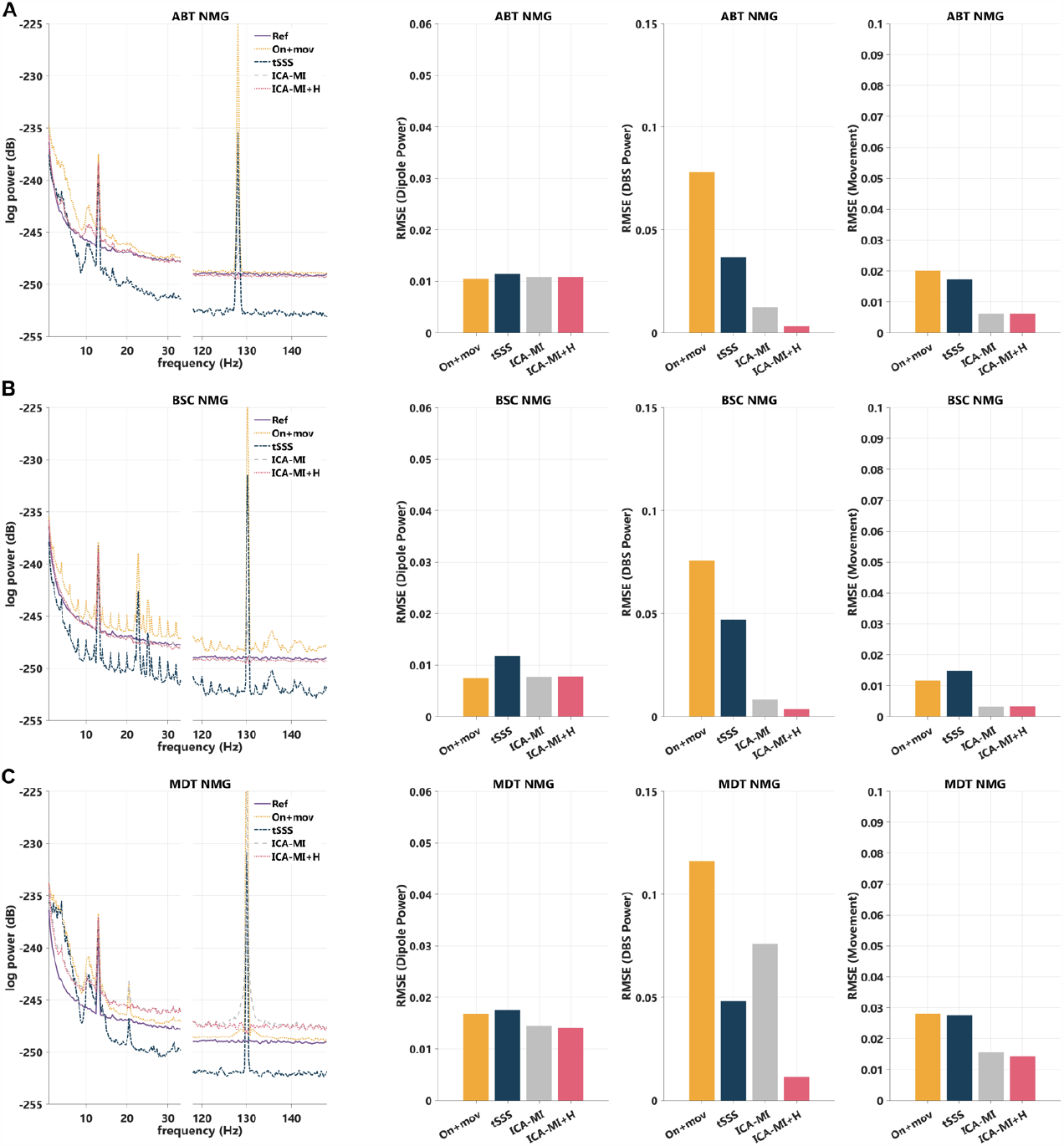
Effective reduction of movement-related and stimulation-induced artefacts in Neuromag Recordings. The first column shows the average power spectrum density (PSD) plots across channels (gradiometers) for Neuromag (NMG) recordings for the three DBS devices from Abbott (ABT), Boston Scientific (BSC) and Medtronic (MDT). Reference recording (Ref) was performed without an implantable pulse generator (IPG) and only the dipole turned on. The three last columns show the RMSE in relation to the reference recording for dipole power (first column), DBS power (second column) and movement (third column). On+mov= dipole, movement and IPG turned on. ICA-MI+H= ICA and mutual information + Hampel filter.

**Figure 5.**
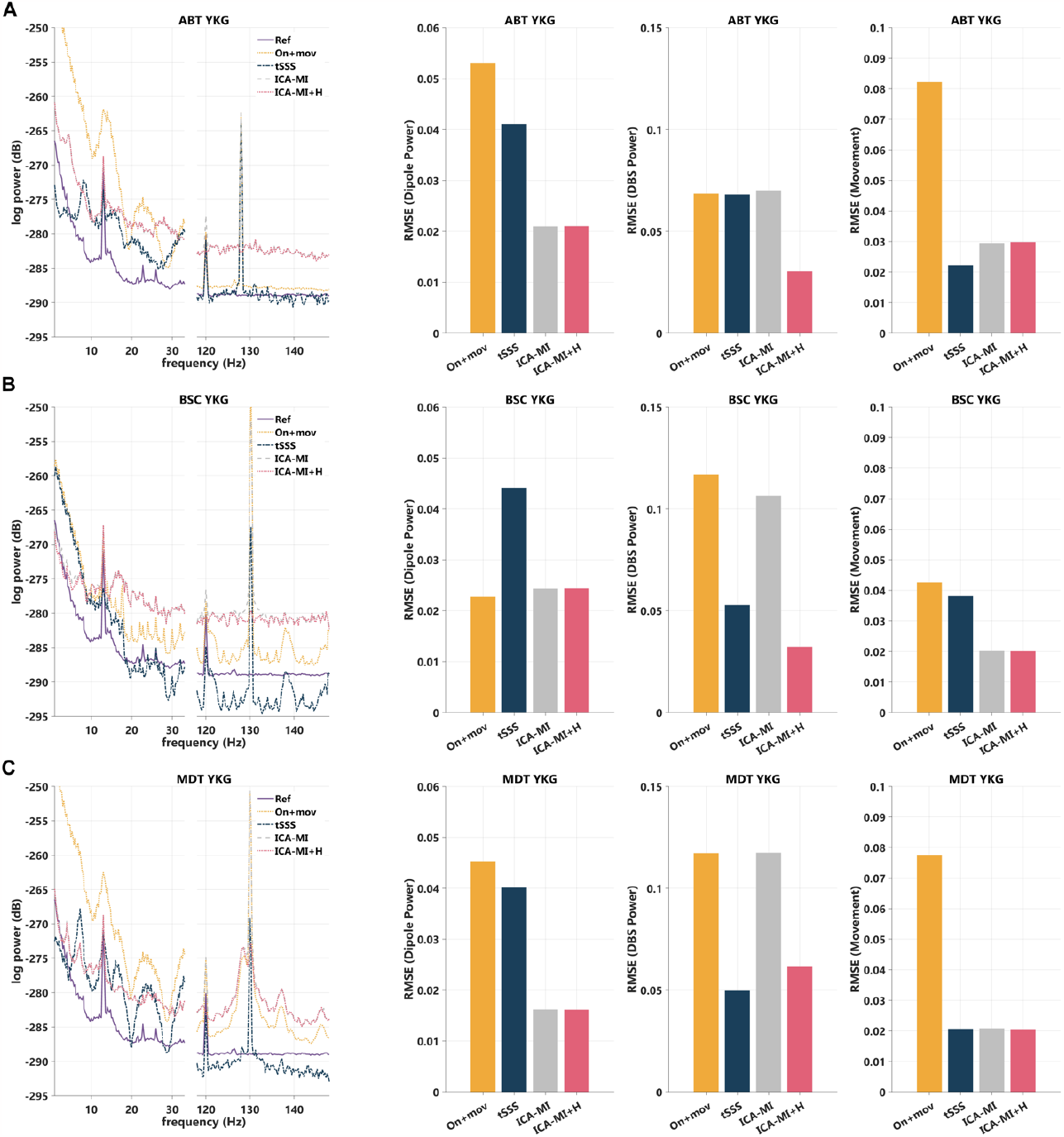
Effective reduction of movement-related and stimulation-induced artefacts in Yokogawa recordings. Legend as in figure 4

### Removal of the DBS artefact

We also compared the performance of established DBS artefact reduction methods on the artefacts described above (figure 4 NMG system and in figure 5 YKG system).

First, dipole activity was largely unchanged by the artefact reduction methods, which indicates that the signal of interest was not reduced by any of the methods. In the YKG recordings, movement related artefacts partially obscured the dipole activity (Fig. 5 A, C), which leads to higher RMSE values in the On+mov condition.

Second, tSSS shifted the baseline downwards, especially in the NMG recordings (Fig. 4). Therefore, tSSS results have to be interpreted differently and the performance as reflected by the RMSE graphs is worse than depicted in figure 4. In fact, besides shifting the baseline, tSSS did not alter the signal.

Third, the ICA-MI approach performed well, especially in reducing the movement-related artefact as well as the electrical stimulation artefact in some of the recordings (Fig. 4, 5). When combining the Hampel filter and ICA-MI, both the movement and electrical stimulation artefact were reduced. In the YKG recordings, ICA-MI led to an upward baseline shift, which means that – in contrast to the tSSS baseline shift – the artefact reduction performance is better than shown in the RMSE results (Fig. 5). Furthermore, ICA-MI alone reduced the IPG-related artefacts in the case of BSC recordings (Fig. 4 B, 5 B). In the stimulation OFF condition IPG-related artefacts were also reduced with ICA-MI for BSC recordings, while for ABT and MDT recordings, IPG-related artefacts were reduced to a smaller extent (Supplementary Fig. 3).

Finally, movement-related artefacts induce low-frequency changes that resemble physiological alpha activity, especially in the case of NMG recordings (Fig. 4). When applying artefact reduction methods, there are residuals of this artefactual alpha peak. For the YKG recordings, the reduction of the movement artefact also leads to low-frequency peaks in the spectrum (Fig. 5).

## Discussion

In this study, we characterize device-specific DBS artefacts in MEG recordings using a phantom setup. We identified IPG-related artefacts, contaminating the power spectrum across various physiologically relevant frequencies. Importantly, even when turned off, all devices generate electrical pulses, which lead to specific additional artefacts in the stimulation OFF condition. We show that movement-related artefacts differ across DBS devices and MEG systems with weaker artefacts for NMG recordings with BSC devices. The electrical stimulation induced artefacts at the stimulation frequency and at lower frequencies. The reduction of stimulation, movement, and IPG-related artefacts was most successful when combining ICA-MI and Hampel filter across recordings and DBS devices. However, especially in ABT and MDT recordings the cleaning of movement-related artefacts resulted in artefactual spectral peaks within physiologically relevant frequencies below 20 Hz.

We observed IPG-related artefacts, that occur even when the IPG is switched off. Naturally, when studying the effect of DBS, a comparison between DBS on and off conditions is used. However, especially when IPG-related artefacts only occur in one of the two conditions, the comparison might be flawed and for instance a reduction of alpha power due to DBS might be a result of decreased IPG or movement-related artefacts, which can mimic physiological activity after artefact reduction. In fact, when cleaning IPG-related artefacts in the stimulation OFF condition for ABT and MDT recordings, the low-frequency power is larger after ICA-MI (Supplementary Fig. 3 D, F) than in the stimulation On recording (Fig. 5 B, C).

The aforementioned movement artefacts relate to the ferromagnetic properties of the DBS hardware components (Bahners et al., 2020; Kandemir et al., 2020). In comparison to ABT and MDT devices, the BSC hardware induced the weakest movement-related artefacts. In the meantime, during our study, MDT developed new extension cables with less ferromagnetic components. Continuous follow-up studies with new devices are needed to assess the impact on MEG artefacts.

The electrical stimulation induced artefacts were consistent across DBS devices, except for the stimulation frequency, which was lower for the ABT device and has been reported before (Kandemir et al., 2020). Furthermore, we observed peaks across different lower frequencies that probably represent sub-harmonics due to the under-sampling of the DBS pulse (Jech et al., 2006; Oppenheim et al., 1997; Sun et al., 2014). These artefacts can be effectively reduced using the Hampel filter as has been demonstrated before (Kandemir et al., 2020). ICA-MI by itself also reduced the electrical artefact to some degree (Fig. 4, 5). Unlike previous studies, we used the channel with the strongest movement-related artefact as a reference for the MI algorithm and not the stimulation artefact itself. This might explain why the algorithm does not reduce the electrical artefact to a larger extent. Instead of using a reference EMG recording above the IPG, we decided to use the channel with the largest movement-related artefact as reference for the MI algorithm, because the movement of the DBS system and particularly the movement of extension cables is not directly reflected in the EMG signal (Abbasi et al., 2016). Yet, the use of a MEG channel with high amplitude low-frequency activity (i.e. movement-related artefacts) as reference for ICA-MI might result in a rejection of physiological low-frequency activity as well. At least in our experimental setup, ICA-MI did not reduce the simulated narrow-band brain signal. Finally, with a combination of both, ICA-MI and Hampel filter, the reduction of stimulation- and movement-related artefacts was most successful and even supressed the aforementioned IPG-related artefacts. In a supplementary patient analysis we observed no reduction in alpha power employing the ICA-MI algorithm with a MEG sensor as reference signal as described above (Supplementary Fig. 4 A). Furthermore, the IPG-related artefact appears to be larger than the electrical stimulation artefact in the time series in this patient example (Supplementary Fig. 4 C). Also in the patient case (BSC device), the IPG-related artefact was reduced.

Surprisingly, tSSS neither reduced the movement-related nor the stimulation-induced artefacts. There are two main differences between our study and the work conducted before, that might explain the poor performance of tSSS (Kandemir et al., 2020). First, the movement-related artefact was weaker in our study. Second, the phantom head was slightly smaller (i.e. slimmer) than the phantom in Kandemir et al, while the cables were attached on top of the head in our study and were in similar proximity to the MEG sensors as in Kandemir et al. (2020). These two factors may have led to a poor performance of tSSS, because the algorithm might not have been able to differentiate the artefact from inner sphere activity.

We also observed MEG system-specific differences in artefacts and overall noise level. First, the frequency range and appearance of movement-related artefacts differed between MEG systems. In the NMG recordings, movement-related artefacts resulted in increased low-frequency power and an artefactual alpha peak, while for the YKG system, the artefactual activity spans to the beta band (Fig. 2 A, C, D, F). Second, the YKG system generally seems to be more sensitive to movement-related artefacts: the RMSE between Reference recording and On+mov recording were larger than for NMG recordings. Third, the previously observed downward baseline shift introduced by tSSS seems to be specific for NMG recordings (Kandemir et al., 2020), while the upward shift due to ICA-MI in the YKG recordings is a new observation. Finally, there were differences between artefact sensitivity of MEG sensors across systems. This is in line with previous work comparing NMG and CTF MEG systems (Kandemir et al., 2020). CTF and YKG share the same type of sensors, namely axial gradiometers with a baseline of 50 mm, while NMG uses planar gradiometers with a baseline of 17 mm. Due to the shorter baseline, the NMG gradiometers suppress distant interfering sources better. These differences in sensor design might account for some of the differences between the systems described above – especially because all other factors (i.e. movement mechanism, phantom head, DBS devices and stimulation parameters) were kept constant for the two recording sessions.

One important limitation of our work is the focus on the sensor space. When localizing the data to source space, the pre-processing pipeline might have to be adjusted depending on the source localization method employed (Jaiswal et al., 2020). In previous work the combination of tSSS and LCMV beamforming revealed the best spatial source mapping in the phantom recording (Kandemir et al., 2020). Importantly, data preprocessed with tSSS needs to be adjusted before applying a beamformer (Jaiswal et al., 2020).

Additionally, the application of LCMV beamforming was sufficient to reduce the movement-related artefacts in studies, which recruited patients in the perioperative stage (i.e. with externalized cables and an external pulse generator) (Oswal et al., 2016a, 2016b). Meanwhile a study in patients with implanted DBS systems found that the artifact reduction using LCMV beamforming was not sufficient, especially for the sensors above the extension cables (Bahners et al., 2020). Finally, our work represents an experimental setting, which will not necessarily be transferable to patient recordings in every case. The preprocessing pipeline used for data analysis has to be tailored based on the respective recordings and the amount of artefact contamination.

## Conclusions

In our study, we show that device-specific IPG-related artefacts have to be considered for MEG-DBS studies and can be reduced using standard artefact reduction methods. Movement-related artefacts were weakest for NMG recordings with BSC devices, while the IPG-related artefacts were also strongest for this condition, but could be well controlled by the use of ICA-MI. Another relevant aspect that has to be considered are the remnants of movement-related artefacts after cleaning, which resemble physiological activity. Finally, we show that combined MEG-DBS recordings are feasible in Yokogawa-MEG systems.

## Acknowledgements

We thank Georg Bahners for his help with building the phantom and Pia Hartmann and Luisa Spallek for their assistance during recordings. We also thank Ahmet L. Kandemir who developed the movement-mechanism and provided his scripts for the analysis from the previous publication. We thank Wolf-Julian Neumann for his valuable feedback.

## References

Abbasi, O., Hirschmann, J., Schmitz, G., Schnitzler, A., Butz, M., 2016. Rejecting deep brain stimulation artefacts from MEG data using ICA and mutual information. J. Neurosci. Methods 268, 131–141. 10.1016/j.jneumeth.2016.04.010

Allen, D.P., 2009. A frequency domain Hampel filter for blind rejection of sinusoidal interference from electromyograms. J. Neurosci. Methods 177, 303–310. 10.1016/J.JNEUMETH.2008.10.019

Bahners, B.H., Florin, E., Rohrhuber, J., Krause, H., Hirschmann, J., van de Vijver, R., Schnitzler, A., Butz, M., 2020. Deep brain stimulation does not modulate auditory-motor integration of speech in Parkinson’s disease. Front. Neurol. 11, 655. 10.3389/fneur.2020.00655

Bahners, B.H., Spooner, R.K., Hartmann, C.J., Schnitzler, A., Florin, E., 2023. Subthalamic stimulation evoked cortical responses relate to motor performance in Parkinson’s disease. Brain Stimul. 16, 561–563. 10.1016/J.BRS.2023.02.014

Benabid, A.L., Pollak, P., Gross, C., Hoffmann, D., Benazzouz, A., Gao, D.M., Laurent, A., Gentil, M., Perret, J., 1994. Acute and long-term effects of subthalamic nucleus stimulation of Parkinson’s disease, in: Stereotactic and Functional Neurosurgery. 10.1159/000098600

Chiken, S., Nambu, A., 2016. Mechanism of deep brain stimulation: inhibition, excitation, or disruption ? Neurosci. 22, 313–322. 10.1177/1073858415581986

Deuschl, G., Schade-Brittinger, C., Krack, P., Volkmann, J., Schäfer, H., Bötzel, K., Daniels, C., Deutschländer, A., al., et, 2006. A randomized trial of deep-brain stimulation for Parkinson’s disease. N.Eng.J. Med. 355, 896–908. 10.1056/NEJMoa060281

Harmsen, I.E., Rowland, N.C., Wennberg, R.A., Lozano, A.M., 2018. Characterizing the effects of deep brain stimulation with magnetoencephalography: A review. Brain Stimul. 10.1016/j.brs.2017.12.016

Jaiswal, A., Nenonen, J., Stenroos, M., Gramfort, A., Dalal, S.S., Westner, B.U., Litvak, V., Mosher, J.C., Schoffelen, J.M., Witton, C., Oostenveld, R., Parkkonen, L., 2020. Comparison of beamformer implementations for MEG source localization. NeuroImage 216, 116797. 10.1016/J.NEUROIMAGE.2020.116797

Jech, R., Růžička, E., Urgošík, D., Serranová, T., Volfová, M., Nováková, O., Roth, J., Dušek, P., Mečíř, P., 2006. Deep brain stimulation of the subthalamic nucleus affects resting EEG and visual evoked potentials in Parkinson’s disease. Clin. Neurophysiol. 117, 1017–1028. 10.1016/J.CLINPH.2006.01.009

Kandemir, A.L., Litvak, V., Florin, E., 2020. The comparative performance of DBS artefact rejection methods for MEG recordings. NeuroImage 219, 117057. 10.1016/j.neuroimage.2020.117057

Krack, P., Volkmann, J., Tinkhauser, G., Deuschl, G., 2019. Deep brain stimulation in movement disorders: from experimental surgery to evidence-based therapy. Mov. Disord. 34, 1795–1810. 10.1002/mds.27860

Limousin, P., Brown, P., Marsden, J., Defebvre, L., Rothwell, J., 1998a. Evoked potentials from subthalamic nucleus, internal pallidum and thalamic stimulation in Parkinsonian and postural tremor patients. J. Physiol. 509P, 176P–177P. 10.1111/j.1469-7793.1998.tb00048.x

Limousin, P., Krack, P., Pollak, P., Benazzouz, A., Ardouin, C., Hoffmann, D., Benabid, A.-L., 1998b. Electrical stimulation of the subthalamic nucleus in advanced Parkinson’s Disease. N.Eng.J. Med. 339, 1105–1111. 10.1056/NEJM199810153391603

Litvak, V., Eusebio, A., Jha, A., Oostenveld, R., Barnes, G.R., Penny, W.D., Zrinzo, L., Hariz, M.I., Limousin, P., Friston, K.J., Brown, P., 2010. Optimized beamforming for simultaneous MEG and intracranial local field potential recordings in deep brain stimulation patients. NeuroImage 50, 1578–1588. 10.1016/j.neuroimage.2009.12.115

Litvak, V., Florin, E., Tamás, G., Groppa, S., Muthuraman, M., 2021. EEG and MEG primers for tracking DBS network effects. NeuroImage 224. 10.1016/j.neuroimage.2020.117447

Lozano, A.M., Lipsman, N., Bergman, H., Brown, P., Chabardes, S., Chang, J.W., Matthews, K., McIntyre, C.C., Schlaepfer, T.E., Schulder, M., Temel, Y., Volkmann, J., Krauss, J.K., 2019. Deep brain stimulation: Current challenges and future directions. Nat. Rev. Neurol. 15, 148–160. 10.1038/s41582-018-0128-2

Oppenheim, A. V, Willsky, A.S., Hamid, S., Ding, J.-J., 1997. Linear time invariant systems, in: Signals & Systems. Pearson.

Oswal, A., Beudel, M., Zrinzo, L., Limousin, P., Hariz, M., Foltynie, T., Litvak, V., Brown, P., 2016a. Deep brain stimulation modulates synchrony within spatially and spectrally distinct resting state networks in Parkinson’s disease. Brain 139, 1482–1496. 10.1093/BRAIN/AWW048

Oswal, A., Jha, A., Neal, S., Reid, A., Bradbury, D., Aston, P., Limousin, P., Foltynie, T., Zrinzo, L., Brown, P., Litvak, V., 2016b. Analysis of simultaneous MEG and intracranial LFP recordings during Deep Brain Stimulation: A protocol and experimental validation. J. Neurosci. Methods 261, 29–46. 10.1016/j.jneumeth.2015.11.029

Piitulainen, H., Bourguignon, M., De Tiège, X., Hari, R., Jousmäki, V., 2013. Coherence between magnetoencephalography and hand-action-related acceleration, force, pressure, and electromyogram. NeuroImage 72, 83–90. 10.1016/J.NEUROIMAGE.2013.01.029

Spooner, R.K., Bahners, B.H., Schnitzler, A., Florin, E., 2023. DBS-evoked cortical responses index optimal contact orientations and motor outcomes in Parkinson’s disease. npj Park. Dis. 9, 1–11. 10.1038/s41531-023-00474-4

Sun, Y., Farzan, F., Dominguez, L.G., Barr, M.S., Giacobbe, P., Lozano, A.M., Wong, W., Daskalakis, Z.J., 2014. A novel method for removal of deep brain stimulation artifact from electroencephalography. J. Neurosci. Methods 237, 33–40. 10.1016/J.JNEUMETH.2014.09.002

Tadel, F., Baillet, S., Mosher, J.C., Pantazis, D., Leahy, R.M., 2011. Brainstorm: A user-friendly application for MEG/EEG analysis. Comput. Intell. Neurosci. 2011. 10.1155/2011/879716

Taulu, S., Hari, R., 2009. Removal of magnetoencephalographic artifacts with temporal signal-space separation: demonstration with single-trial auditory-evoked responses. Hum. Brain Mapp. 30, 1524–1534. 10.1002/hbm.20627

Welch, P.D., 1967. The Use of Fast Fourier Transform for the Estimation of Power Spectra: A Method Based on Time Averaging Over Short, Modified Periodograms. IEEE Trans. Audio Electroacoust. 15, 70–73. 10.1109/TAU.1967.1161901

