## SupplementaryData for "Deep brain stimulation device-specific artefacts in MEG recordings"

### Supplementary Data

In both MEG systems data was sampled at 5,000 Hz with a high-pass of 0.1 Hz and a low-pass of 1,660 Hz. In addition to phantom recordings, a patient recording was performed in Berlin for supplementary analysis with the same acquisition settings as for the phantom recording. The patient had a Boston Scientific (BSC) device with pallidal DBS electrodes and was stimulated with a bipolar montage and a stimulation frequency of 185 Hz. The patient gave their prior written informed consent, the study was approved by the local ethics committee at Charité Berlin (proposal number: EA2/035/20), and performed in accordance with the Declaration of Helsinki (WMA, 2013).

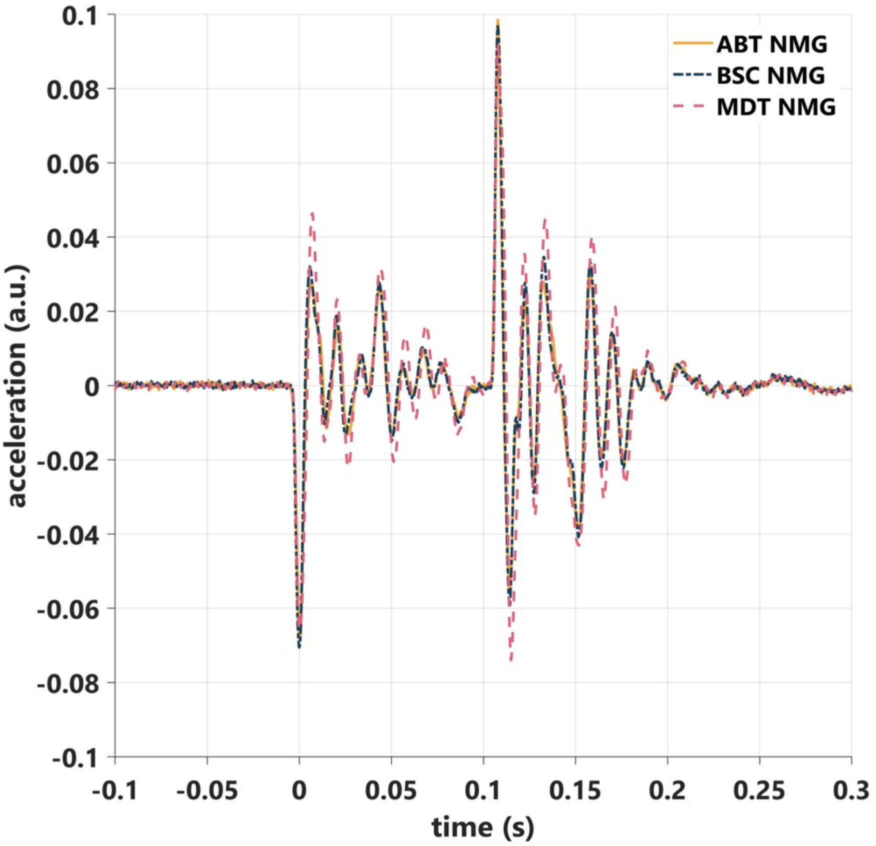


**Supplementary Figure 1. Movement-related artefacts did not differ across experimental conditions with different DBS devices.** Average acceleration across movement events (n=21) for each of the different DBS devices and respective recordings (ABT= Abbott, BSC= Boston Scientific, MDT: Medtronic). Time point 0 s marks the onset of movement.

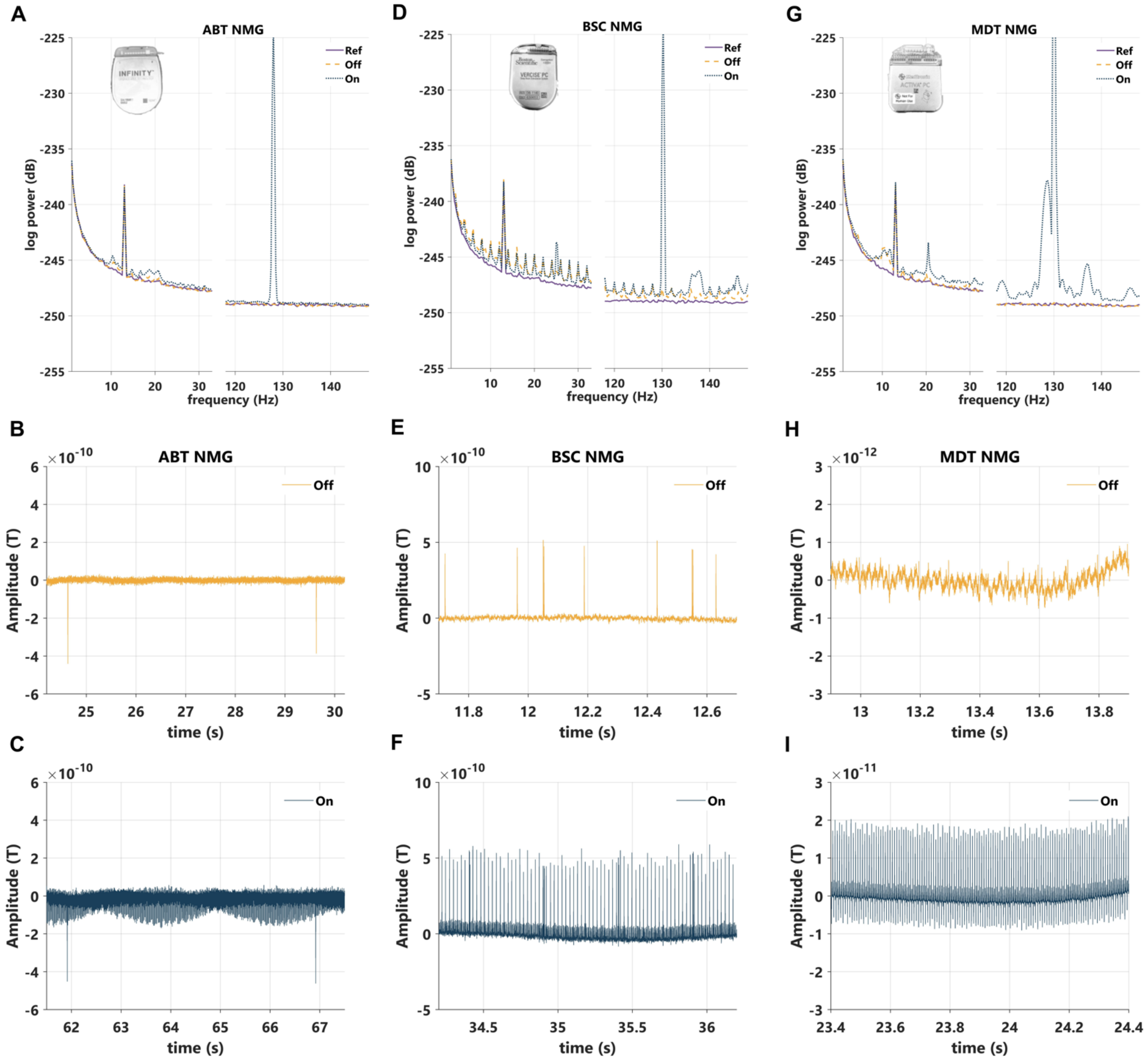


**Supplementary Figure 2. IPG-related Artefacts influence signal quality in frequency and time domain in Neuromag recordings. A, D, G** Average power spectrum density (PSD) plots across channels (gradiometers) for Neuromag (NMG) recordings with DBS devices from Abbott (ABT), Boston Scientific (BSC) and Medtronic (MDT). Reference recording (Ref) was performed without an implantable pulse generator (IPG) and only the dipole turned on. The Off condition was recorded with the IPG inside the magnetic-shielded room and the IPG turned Off, while only dipole activity was turned on as in the reference file. The On condition was recorded with the IPG turned On and only the dipole activity turned on without movement. **B, C, E, F, H, I**. Exemplary time series signal showing the IPG-related artefacts in the stimulation Off (B, E, H) and On (C, F, I) conditions. Note that for MDT recordings the artefact could only be observed in magnetometers, which is why a magnetometer time series is shown.

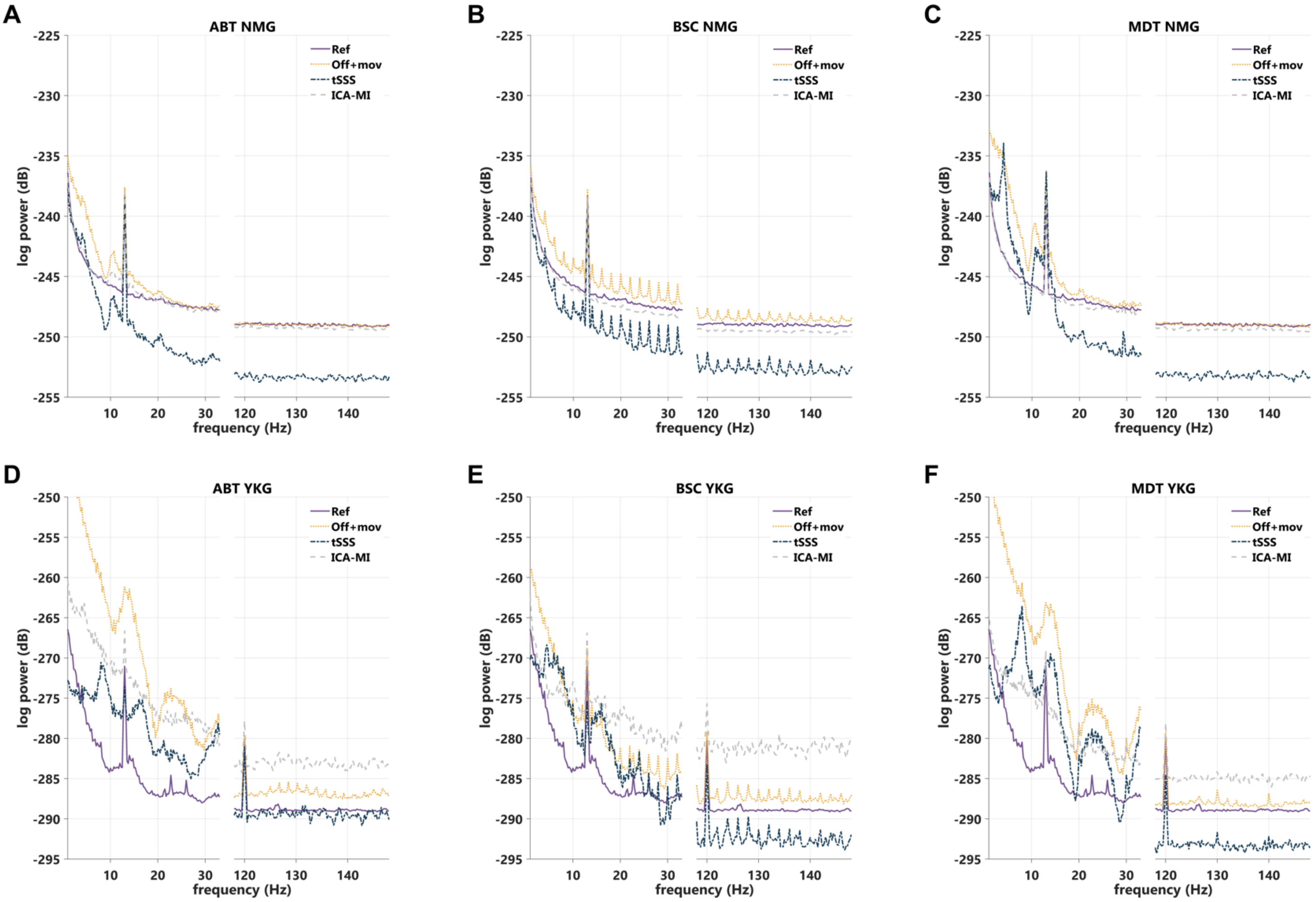


**Supplementary Figure 3. Reduction of movement-related and IPG-related artefacts in the stimulation OFF condition for Yokogawa and Neuromag recordings.** Average power spectrum density (PSD) plots across channels (gradiometers) for Neuromag (NMG) and Yokogawa (YKG) recordings (NMG: A-C, YKG: D-F). The PSD for the three DBS devices from Abbott (ABT) are shown in A and D, from Boston Scientific (BSC) in B and E and from Medtronic (MDT) in C and F. Reference recording (Ref) was performed without an implantable pulse generator (IPG) and only the dipole turned on. Off+mov= dipole, movement, wires and IPG turned off.

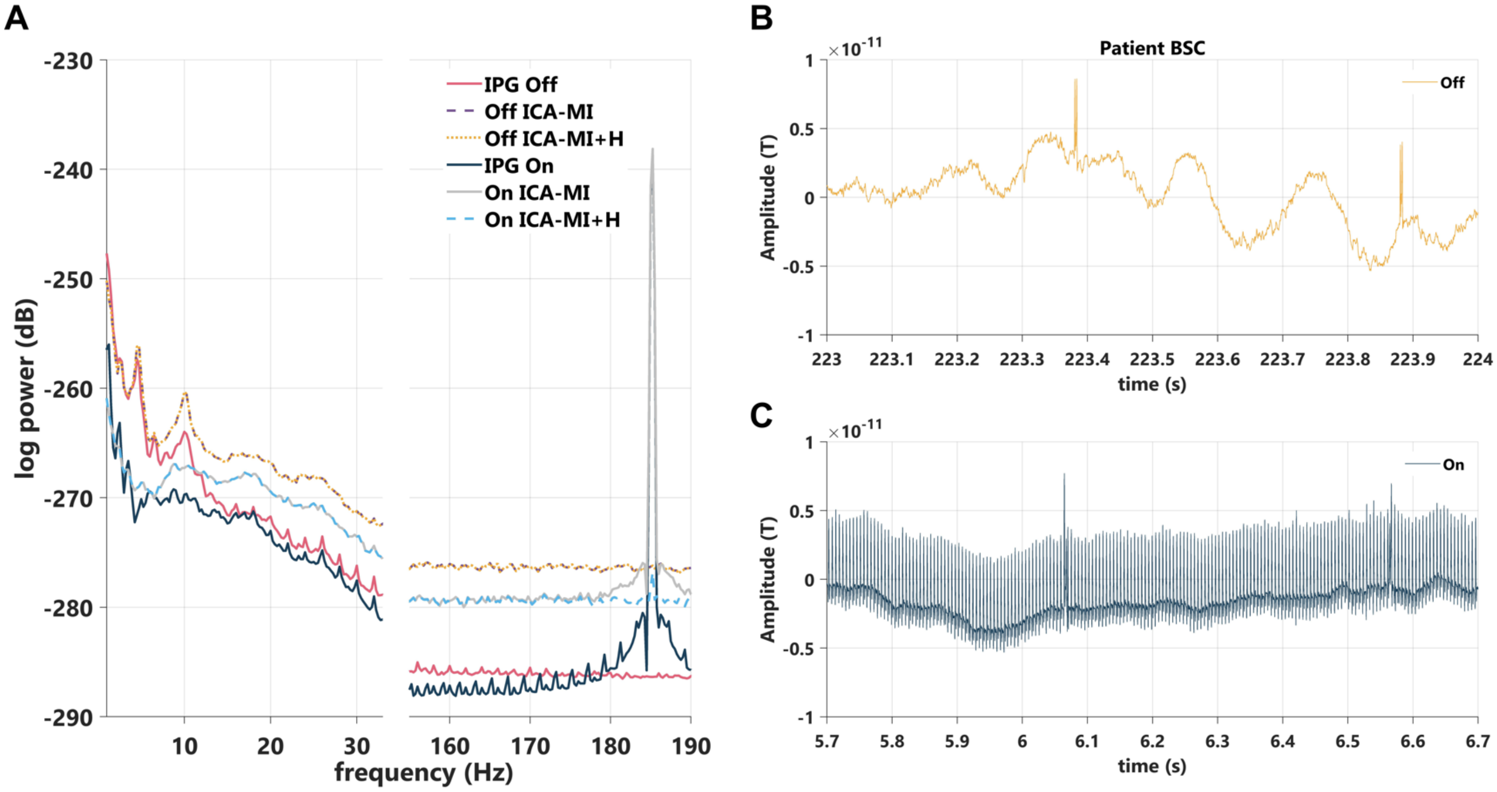


**Supplementary Figure 4. Movement-related and IPG-related artefacts and their reduction in the stimulation OFF and ON condition for Yokogawa recordings in a patient with a BSC system. A.** Average power spectrum density (PSD) plot across channels. ICA-MI and Hampel filter were used for both, the recording with the IPG Off and On. **B, C** Sensor time series for representative left (side of IPG implantation) temporal channel in the stimulation Off (B) and On (C) conditions. The IPG-related peaks in the time series are larger than the stimulation artefact and can be seen across the spectrum before cleaning as peaks with a distance of 2 Hz in the IPG Off and IPG On conditions (A).

### References

WMA, 2013. World medical association declaration of Helsinki: Ethical principles for medical research involving human subjects. JAMA 310, 2191–2194. https://doi.org/10.1001/JAMA.2013.281053
